# Plasmid Profiler: Comparative Analysis of Plasmid Content in WGS Data

**DOI:** 10.1101/121350

**Authors:** Adrian Zetner, Jennifer Cabral, Laura Mataseje, Natalie C Knox, Philip Mabon, Michael Mulvey, Gary Van Domselaar

## Abstract

**Summary:** Comparative analysis of bacterial plasmids from whole genome sequence (WGS) data generated from short read sequencing is challenging. This is due to the difficulty in identifying contigs harbouring plasmid sequence data, and further difficulty in assembling such contigs into a full plasmid. As such, few software programs and bioinformatics pipelines exist to perform comprehensive comparative analyses of plasmids within and amongst sequenced isolates. To address this gap, we have developed Plasmid Profiler, a pipeline to perform comparative plasmid content analysis without the need for *de novo* assembly. The pipeline is designed to rapidly identify plasmid sequences by mapping reads to a plasmid reference sequence database. Predicted plasmid sequences are then annotated with their incompatibility group, if known. The pipeline allows users to query plasmids for genes or regions of interest and visualize results as an interactive heat map.

**Availability and Implementation:** Plasmid Profiler is freely available software released under the Apache 2.0 open source software license. A stand-alone version of the entire Plasmid Profiler pipeline is available as a Docker container at https://hub.docker.com/r/phacnml/plasmidprofiler_0_1_6/.

The conda recipe for the Plasmid R package is available at: https://anaconda.org/bioconda/r-plasmidprofiler

The custom Plasmid Profiler R package is also available as a CRAN package at https://cran.r-project.org/web/packages/Plasmidprofiler/index.html

Galaxy tools associated with the pipeline are available as a Galaxy tool suite at https://toolshed.g2.bx.psu.edu/repository?repository_id=55e082200d16a504

The source code is available at: https://github.com/phac-nml/plasmidprofiler

The Galaxy implementation is available at: https://github.com/phac-nml/plasmidprofiler-galaxy

Address: National Microbiology Laboratory, Public Health Agency of Canada, 1015 Arlington Street, Winnipeg, Manitoba, Canada

**Supplementary information:** Documentation: http://plasmid-profiler.readthedocs.io/en/latest/

## 1. Introduction

Plasmids are circular or linear extrachromosomal double-stranded DNA molecules capable of self-replication in a host and transfer between host cells. They are highly variable both in length (from one to several hundred kilobases) and number of copies per cell with many plasmids harboring genes beneficial to host fitness and pathogenicity. These genes confer benefits like antimicrobial resistance (AMR), organic product degradation, and virulence factors such as toxin production. Carriage and transfer of pathogenicity and AMR determinants makes development of epidemiologically relevant tools and plasmid classification systems important health care research topics. It was to these ends that in response to a multi-species plasmid-mediated drug-resistant outbreak Plasmid Profiler was developed to explore and visualize cellular plasmid content directly from whole genome sequencing short reads.

The backbone of a given plasmid contains genes essential to its maintenance and stable inheritance (Nordstrom, K. and Austin, S.J. 1989, Shintani, M., Sanchez, Z.K., et al. 2015). This gives rise to one common method of plasmid classification driven by the concept of incompatibly grouping where two plasmids with the same incompatibility group cannot replicate within the same cell. These groups are defined by the plasmids replication or equipartitioning systems (Couturier, M., Bex, F., et al. 1988). Currently, we are able to screen for 116 incompatibility groups and subgroups based on the sequences within the Enterobacteriaceae plasmid typing database provided by PlasmidFinder (Carattoli, A., Zankari, E., et al. 2014).

The role of plasmids in health and disease makes them important to characterize; however, due to their structure it is stubbornly difficult to do so. Extensive mosaicism, repetitive sequences, and smaller mobile elements (eg. transposons) within plasmids collectively impede the ability to discriminate between plasmid and chromosomal DNA in whole genome sequencing (WGS) data analysis.

Without resorting to long-read sequencing, assembling a full plasmid sequence from short reads is difficult and success not guaranteed even when used in conjunction with mate-pair technologies (Lynch, T., Petkau, A., et al. 2016). Some software packages have made attempts to address plasmid analysis from short read sequencing data including PLACNET (Lanza, V.F., de Toro, M., et al. 2014), Recycler (Rozov, R., Brown Kav, A., et al. 2016), plasmidSPAdes (Antipov, D., Hartwick, N., et al. 2016), PlasmidFinder (Carattoli, A., Zankari, E., et al. 2014), but regretfully to date no system has been shown to be capable of reliably reconstructing plasmids (Arredondo-Alonso, S., van Schaik, W., et al. 2017, Orlek, A., Stoesser, N., et al. 2017). Accordingly, there is a scarcity of software available to explore the unassembled plasmid content of bacterial short-read WGS data. We have developed the Plasmid Profiler pipeline to explore plasmid content without having to resort to the time-consuming and unreliable *de novo* assembly process.

## 2. Methods

Plasmid Profiler is implemented as a Galaxy workflow and operates as follows:

**1. Three inputs are required: WGS short reads in FastQ format, the supplied plasmid sequence database, and the supplied but user-modifiable replicon sequence and gene of interest database**. The plasmid sequence database originates as a multi-fasta file from NCBI starting with all Gammaproteobacteria plasmid sequences (n=2797, size range: 1065-727,905 bp). Database curation was performed by first removing redundancy using CD-HIT (v.4.6) (Fu, L., Niu, B., et al. 2012, Li, W. and Godzik, A. 2006) to cluster sequences at >99% sequence identity and >99% length, then identify a representative sequence from each cluster. All sequences containing the following annotation keywords: “draft”, “contig”, “scaffold”, “partial”, “incomplete”, and “putative” were removed to avoid incomplete and low quality plasmid sequence data. The database was then formatted for SRST2 using the Python scripts included in the SRST2 distribution. The user-modifiable replicon and genes of interest database is a multi-fasta file with plasmid replicon sequences from PlasmidFinder (Center for Genomic Epidemiology’s PlasmidFinder database available at: https://cge.cbs.dtu.dk//services/data.php) along with user-defined nucleotide sequences for genes of interest. These three elements are then passed to the tools of the workflow.

**2. KAT (v. 2.3.1) (Mapleson, D., Garcia Accinelli, G., et al. 2017) removes unrepresented Gammaproteobacteria plasmid sequences to create an individualized plasmid database per sample**. As a measure to increase speed and efficiency, a customized database is created for each WGS isolate to contain only sequences moderately homologous to the isolate’s sequence data. This per isolate database reduction step is accomplished using KAT to compare the k-mers of each isolate’s WGS reads to a series of k-mer hashes derived from the entire Gammaproteobacteria plasmid database. Database sequences which are represented at a minimum of 80% by the k-mers of a particular WGS isolate are retained in that isolate’s individual plasmid database and all other sequences discarded. This process rapidly creates individual databases for each WGS sample and avoids the memory intensive and time consuming mapping process to references unrepresented in an isolate’s WGS reads.

**3. SRST2 (v.0.1.6) (Inouye, M., Dashnow, H., et al. 2014) identifies putative plasmid hits from the individual databases**. SRST2 is a software package that allows for *in silico* sequence typing of WGS data based on the mapping of short sequence reads to a custom gene and/or allele database. Paired or single end reads are mapped with Bowtie2 (Langmead, B. and Salzberg, S.L. 2012) and the resulting pileups scored according to sequence coverage and divergence. Plasmid Profiler replaces the SRST2 gene database with the curated plasmid database from Step 1. At this stage of the pipeline, SRST2 is run using the “Custom Virulence Database” parameter with the individualized plasmid databases serving as the SRST2 database for their respective isolate.

**4. BLAST+ (v 2.2.31) (Camacho, C., Coulouris, G., et al. 2009) identifies the incompatibility groups and genes of interest found in the high scoring sequences from step 3**. Plasmid sequences identified with SRST2 are used to create a custom BLAST database against which 116 plasmid replicon sequences (Center for Genomic Epidemiology’s PlasmidFinder database available at: https://cge.cbs.dtu.dk//services/data.php) are queried using MegaBLAST (Morgulis, A., Coulouris, G., et al. 2008). Included in the query were the coding sequences for user-specified genes of interest (eg. AMR genes). The query sequences (PlasmidFinder replicons + AMR) are presented in a multi-FASTA file with AMR genes or genes of interest appended to the end, prefixed with the tag “(AMR)”. Five carbapenemase genes are included in the default database as examples to be modified or replaced as the user decides. Only the best hit per plasmid is displayed on the final heat map.

**5. The Plasmid Profiler R package parses and visualizes the results of steps 2-4**. The following steps are completed using a series of functions written in the R programming language and published in the Plasmid Profiler R package which is available through CRAN. The percent ID (PID) of each BLAST hit is adjusted by multiplying it by the ratio of query to hit length to compensate for hits with high PID but incomplete sequence coverage. Hits of less than 80% adjusted PID are discarded. Each identified plasmid is labelled with its associated incompatibility group (if no match is found then this is reported as “-”) and presence/absence of the user-specified genes of interest (eg. AMR genes). Sureness values are calculated for each sample from the sum of sequence coverage (normalized over the range 0 to 1) and sequence divergence (normalized over the range -1 to 0). This measure is then normalized over the range 0 to 1. The Sureness values are used to create a distance matrix upon which hierarchical clustering using complete linkage is applied creating a dendrogram and determining the ordering of isolates on the resulting plot. This dendrogram reveals the similarity of putative plasmid content in the samples examined. A tile geometry is plotted using ggplot2 (v2.1.0) (Wickham, H. 2009) to simulate a heatmap with plasmids ordered and coloured by incompatibility group then arranged by decreasing Sureness value with correspondingly decreasing alpha values (Fig. 1). This heatmap is reproduced using plotly (v4.5.6) (Plotly Technologies Inc.) to allow for interaction with the output in a web browser. The post-mapping analysis steps can be run multiple times with varying filters such as length, Sureness value, or mapping coverage applied.

**Figure 1.**
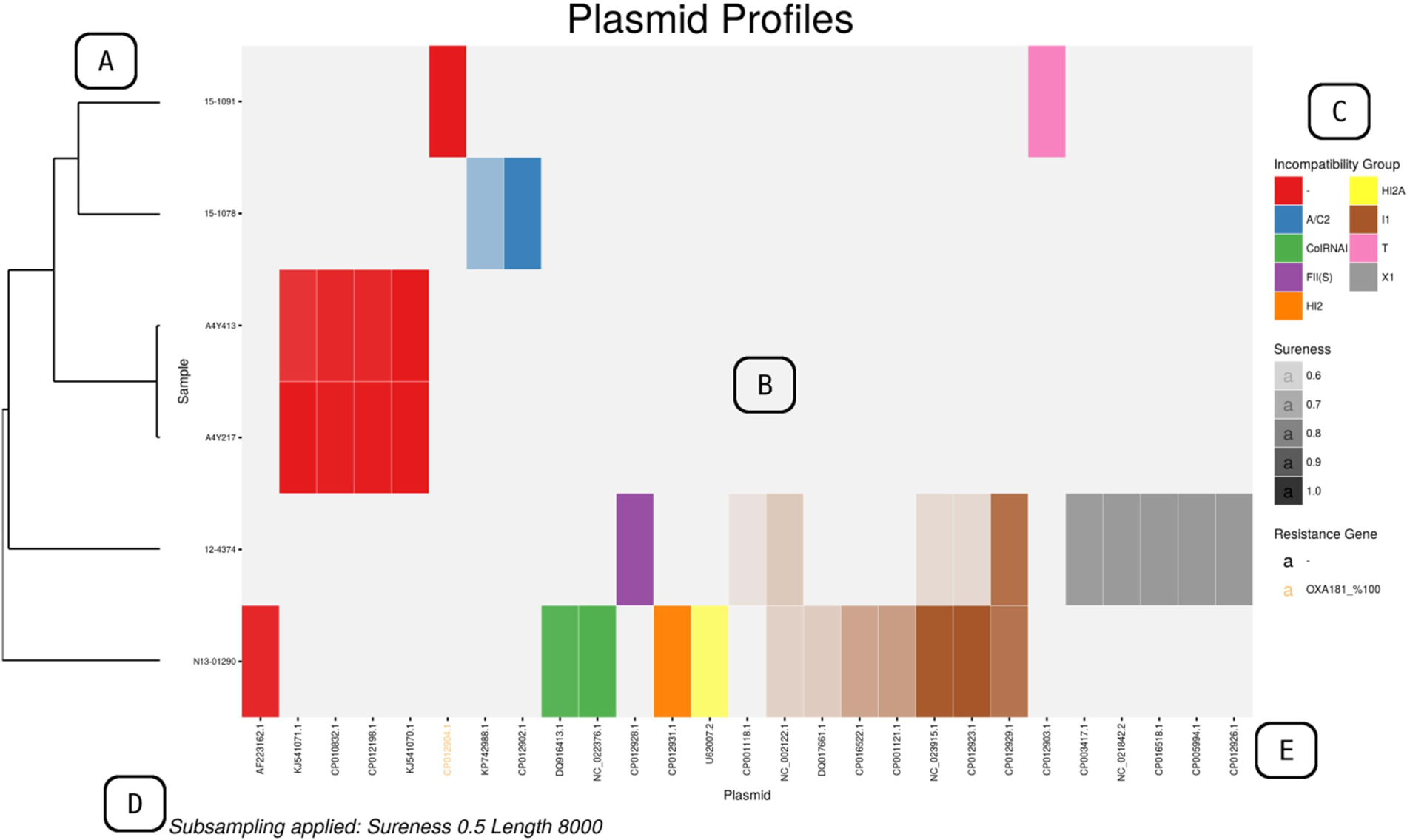
An example PNG output of Plasmid Profiler. A) dendrogram revealing plasmid content similarity amongst samples; B) heatmap created using ggplot2’s tile geometry where colour indicates incompatibility group membership and alpha value the Sureness value over a range from filter cut-off to 1; C) legends for Sureness value, incompatibility group colour guide (“-” if unknown), and AMR gene or user-specified gene of interest colour and identity; D) any data filtering applied is noted here; E) plasmid names below the heatmap are coloured according to the best hit AMR gene or user-specified gene of interest found within that plasmid.

Users can run the pipeline locally through a standalone Galaxy (Afgan, E., Baker, D., et al. 2016) instance in a Docker container (Boettiger, C. 2015). Alternatively, users can install the galaxy pipeline and tools to a pre-existing Galaxy instance from the Galaxy Toolshed (https://toolshed.g2.bx.psu.edu/) (Blankenberg, D., Von Kuster, G., et al. 2014). The R related portion of the analysis is available as a conda recipe and as a package from the Comprehensive R Archive Network (CRAN) that can be easily installed to a local R instance for further manipulation of the mapping output.

## 3. Conclusion

Plasmid Profiler is a useful comparative tool for visualization of plasmid content in WGS data from bacterial isolates without resorting to difficult and uncertain assembly methods or the use of long-read sequencing technologies. It addresses a gap in current plasmid-mediated outbreak analysis methods where an overview of plasmids and antimicrobial resistance genes within a collection of isolates is difficult to obtain. Although the tool is limited by plasmids present in the database, identification and visualization of plasmid backbones within outbreak organisms is important to understanding and conveying the full context of an outbreak.

## Acknowledgement

The authors would like to acknowledge the following people and groups for their help in bringing this software to life: the Bioinformatics Core at the National Microbiology Laboratory (especially Eric Enns, Philip Mabon, and Aaron Petkau), Chand Mangat for providing the necessary PacBio sequence data for pipeline validation, and the Galaxy team for their rapid response to issues raised during development as well as the integration within Galaxy for some of the necessary bioinformatics tools used by Plasmid Profiler.

